# Comparison of immunohistochemistry methods in embryonic chicken corneal tissue

**DOI:** 10.64898/2026.03.30.715369

**Authors:** Joshua T. Harkins, Mitchell M. Hill, Jena L. Chojnowski

## Abstract

Immunohistochemistry (IHC) is widely used to assess protein expression in corneal tissue, yet staining outcomes are strongly influenced by tissue preparation methods and regional differences within the cornea. This study aimed to systematically compare three processing techniques including paraffin (wax) embedding, wax embedding with antigen retrieval (wax AR), and cryosectioning for IHC analysis in embryonic day 18 chicken corneal tissue. Markers representing key biological functions were evaluated, including progenitor activity (PAX6, P40), tissue architecture (actin), and immune surveillance (TAP1, CD68), across central and limbal regions. Cryosectioning consistently produced the most specific staining for nuclear and antigen-sensitive markers. PAX6 and P40 exhibited strong, nuclear-localized expression in the corneal epithelium only under cryo conditions, whereas wax-based methods resulted in reduced specificity and irregular signal distribution. TAP1-positive immune cells were detectable in the limbal stroma exclusively in cryosections, highlighting improved antigen preservation. In contrast, actin staining, was best preserved with wax AR, and provided superior structural clarity and expected expression patterns across corneal layers. CD68 showed minimal or inconsistent staining in corneal tissue across all methods despite positive control validation. These findings demonstrate that optimal IHC outcomes in corneal tissue are marker-dependent and influenced by preparation methods and regional tissue context. Cryosectioning is recommended for detecting nuclear and immune-related antigens, while wax AR is preferable for preserving tissue architecture. This study provides a practical framework for improving reproducibility and interpretation of corneal immunostaining in avian models.

**Highlights:**

- Cryosectioning preserves nuclear and immunity antigens in cornea
- Wax AR improves structural markers such as actin in corneal tissue
- Staining outcomes depend on marker and tissue preparation method
- Cryosectioning enables detection of limbal immune-associated cells (TAP1)
- CD68 shows limited corneal expression despite positive control staining

## 1. Introduction

The cornea is a transparent, avascular tissue that forms the anterior surface of the eye and is essential for both refraction and protection. Its optical function depends on a highly ordered architecture that minimizes light scattering while maintaining mechanical integrity and immune quiescence (Meek et al., 2025; Raut et al, 2025). In addition to serving as a physical barrier, the cornea plays an active role in regulating ocular immune responses, as inflammation or vascularization can severely compromise transparency (Niederkorn, 2002; Prieto-Nevarez et al., 2026). Structurally, the cornea consists of three principal cellular layers: a stratified epithelium, a collagen-rich stroma, and an endothelial monolayer (Meek et al., 2025; Raut et al, 2025). The epithelium provides a renewable protective surface sustained by limbal stem cells located at the corneoscleral junction, further known as the limbal region (Meshko et al., 2026). Beneath this, the stroma comprises most of the corneal thickness and derives its strength and optical properties from densely packed, uniformly arranged collagen fibrils along with keratinocytes (Meek et al., 2025). The endothelium, the deepest layer, maintains stromal hydration through active fluid regulation, preserving corneal clarity (Bonanno, 2012). Together, these layers form a highly specialized tissue necessary for sight acuity and overall sight maintenance.

Another critical aspect of the eye to take into consideration when investigating the cornea is the regional specialization within the tissue. The central cornea is optimized for transparency and refractive precision, characterized by an acellular, avascular environment and tightly organized extracellular matrix (Meek et al., 2025). In contrast, the limbal region supports epithelial stem cell maintenance and is enriched with vasculature, stromal heterogeneity, and immune cell populations (Meshko et al., 2026). This regional heterogeneity directly impacts immunohistochemical (IHC) outcomes: dense stromal collagen in the central cornea can restrict antibody penetration and antigen accessibility, while the increased cellular heterogeneity and immune activity of the limbus can elevate background or nonspecific staining.

IHC remains a powerful approach for mapping protein expression across corneal layers and regions, enabling the study of epithelial differentiation, stromal organization, immune surveillance, and vascular boundaries (Mebratie and Dagnaw, 2024). Wax embedding preserves corneal tissue architecture exceptionally well, maintaining the stratified epithelium, layered stroma, and endothelial monolayer with high fidelity. However, wax embedding can mask protein epitopes through fixation-induced crosslinking, which may limit detection of certain intracellular targets. The addition of antigen retrieval steps can overcome limitations of wax embedding by restoring masked epitopes. Heat or enzymatic retrieval breaks crosslinks formed during fixation, allowing improved antibody access to nuclear proteins. As a result, wax embedding with antigen retrieval (wax AR) is a powerful method for balancing preservation of morphology with reliable detection of intracellular proteins. Cryosectioning avoids chemical fixation and instead preserves antigenicity through rapid freezing, making it ideal for detecting proteins sensitive to fixation. While tissue morphology is less crisp than wax sections, cryosectioning provides superior preservation of delicate epitopes. This method prioritizes antigen detection over structural detail, offering advantages for studies focused on immune surveillance and other antigen-sensitive processes in the cornea. To compare staining outcomes across corneal regions and preparation methods, we selected a set of markers representing key biological features of the eye: progenitor activity, tissue architecture, and immune surveillance. Together, these markers provide a broad framework for assessing both central and limbal specialization, as well as responses to experimental or pathological conditions.

Progenitor activity in the corneal epithelium is reflected by nuclear-expressed transcription factors such as PAX6 and P40. PAX6 and P40 are essential for eye development and epithelial identity, regulating stem cell maintenance at the limbus as well as differentiation in the central cornea (Li et al., 2023; More et al., 2024). Their misregulation has strong associations with congenital eye malformations, such as aniridia, and negative long-term eye health (Di Iorio et al., 2023; More et al., 2024). PAX6 expression is found equally throughout the epithelial cell layers, but it has higher expression levels within the epithelium of the limbal region when compared to the central cornea and can be used as an identifying marker for the limbal region. P40 is expressed in progenitor cells, which can be found in the basal epithelial layer of the limbus and the central cornea but does not show differences in expression between the two regions (Li et al., 2023). Together, these markers distinguish between progenitor-rich limbal tissue and more differentiated central corneal epithelial layers, providing insights into corneal renewal and specialization (Meshko et al., 2026).

A tissue architecture marker was included to capture broad features of corneal structure. Actin, a cytoskeletal protein, highlights cell shape, junctional organization, and migratory dynamics within the corneal epithelium and stroma (Pollard and Cooper, 2009; Wilson, 2020). In the cornea, actin staining outlines the epithelium, stroma, endothelium, and vasculature, making it particularly useful for assessing the preservation of tissue architecture following embedding and sectioning (Pollard and Cooper, 2009; Wilson, 2020). Differences in actin signals can reveal preparation-induced differences such as tissue distortion, cell loss, or disrupted epithelial layering. In this study, actin serves as a biological marker of epithelial and stromal organization and for comparing morphological preservation between central and limbal corneal tissue.

Immune surveillance in the cornea was assessed using markers of antigen processing and macrophage populations. Transporter Associated with Antigen Processing 1 (TAP1) is a core component of the major histocompatibility complex (MHC) class I antigen presentation pathway, responsible for transporting cytosolic peptides into the endoplasmic reticulum for loading onto MHC I molecules (Wearsch and Cresswell, 2008; Neefjes et al., 2011). In the cornea, TAP1 expression reflects intrinsic antigen-processing capacity, which is tightly regulated to maintain immune privilege in the central cornea while supporting heightened immune readiness in the limbal region (Niederkorn, 2002; Wearsch and Cresswell, 2008; Prieto-Nevarez et al., 2026).

Regional differences in TAP1 staining therefore provide insight into how corneal tissues balance immune surveillance with the need to preserve optical clarity. TAP1 serves as a readout of antigen presentation machinery within resident corneal cells, linking epithelial and stromal compartments to local immune competence. Cluster of Differentiation 68 (CD68) labels macrophages, which are more prevalent near the limbus than in the central cornea (Niederkorn, 2002; Hamrah et al., 2003; Prieto-Nevarez et al., 2026). The combined analysis of TAP1 and CD68 provides insight into regional immune differences and the cornea’s capacity to respond to injury or inflammatory stimulus, demonstrating the importance of the limbal niche in coordinating immune surveillance while preserving tissue homeostasis and transparency.

Despite widespread use of IHC in corneal research, few studies have systematically compared how tissue preparation methods influence antibody staining outcomes across distinct corneal regions (Young et al., 2014; Mebratie and Dagnaw, 2024). There is a lack of standardized reference data evaluating how wax embedding, wax AR, and cryosectioning perform in embryonic chicken corneal tissue. This gap limits reproducibility and complicates cross-study comparisons (Mebratie and Dagnaw, 2024). The aim of this study is to directly compare tissue preparation methods: wax embedding, wax AR, and cryosectioning for antibody staining across the diversity of corneal tissue in embryonic chicken eyes. By assessing multiple markers spanning progenitor identity, tissue architecture, and immune surveillance, this work seeks to evaluate staining robustness, regional specificity, and methodological trade-offs. Ultimately, this study provides a practical reference framework for optimizing IHC in avian ocular tissue and supports more reproducible and interpretable corneal immunostaining across laboratories.

## 2. Materials and Supplies

### 2.1 Embryonic chicken embryos

Fertilized chicken eggs were provided by Tyson Farms Complex of Monroe, North Carolina, and stored at 15°C until incubation. Eggs were incubated in a humidity controlled, 35°C rocking incubator until embryonic age 18 (E18) for all experiments; Hamburger and Hamilton stages 44-45 (Hamburger and Hamilton, 1992). Chick embryos were removed from the eggshell and yolk sac and then decapitated, and corneal tissue was dissected from the eye and fixed with 4% paraformaldehyde (PFA) overnight,

## 3. Detailed Methods

### 3.1 Tissue Processing

#### 3.1.1 Wax Embedding and Sectioning

The PFA fixed corneas were washed 3x of 30 min 1x PBS then placed into 70% ethanol overnight before they were dehydrated (increasing gradient of ethanol) and embedded into wax. Wax-embedded tissues were then sectioned at 10 μm using a microtome. Slides were stored at room temperature (RT) before hematoxylin and eosin (H&E) staining or IHC.

#### 3.1.2 Cryo Embedding and Sectioning

The PFA fixed corneas were washed 3x of 30 min 1x PBS then placed into 10% sucrose and 20% sucrose solution washes for 1hr each at 4°C. A final 30% sucrose solution wash was done overnight at 4°C. Samples were then washed 3x of 30 min 1x PBS before being embedded into OCT and frozen at -80°C. Samples were sectioned using a cryostat at 10 μm, and the slides were stored at 4°C. Slides were thawed for one hour at RT before IHC was performed.

#### 3.1.3 Antigen Retrieval

Some wax-embedded sections received an antigen retrieval (AR) step by using a 10 mM sodium citrate buffer (pH 6.0) with 0.05% Tween-20 for 15 minutes at 95°C. Slides rested for 10 minutes while in sodium citrate buffer before allowing to cool down for handling purposes. Slides were then washed 3x of 5 min 1x PBS at RT before IHC was performed.

### 3.2 Histology

#### 3.2.1 H&E

Wax-embedded tissues sectioned onto slides were deparaffinized and rehydrated before being stained with hematoxylin for 5 minutes and rinsed with tap water. They were then stained with eosin for 2 minutes, dehydrated, and cleared with two changes of xylene for 2 minutes each. The slides were mounted and cover-slipped using cytoseal^TM^ 60 before microscopic imaging.

#### 3.2.1 Immunohistochemistry

IHC was performed on wax-embedded, wax AR, and cryo-sectioned samples. All samples were permeabilized with a 1% Triton X-100 solution at RT for 30 minutes. Then samples were blocked in 2% bovine serum albumin (BSA), 2% goat serum, 1% Triton X-100 in 1x Tris Buffered Saline (TBS) at RT for 2 hours. Slides were then incubated with monoclonal primary antibodies Rb P40 (Abcam AB203826, 1:200), Rb Cd68 (Abcam AB213363, 1:100), Rb Pax6 (Abcam AB195045, 1:500), Ms Pax6 (Abcam AB78545, 1:100), Ms CD11c (Abcam AB254183, 1:250), Ms Actin (DSHB AB_528068, 1:8.5), Ms Pax6-S (DSHB AB_528427, 1:12), and Ms TAP1 (DSHB AB_531780, 1:55). Dilutions were either taken from the technical sheet or by titration of different concentrations while comparing staining intensities and specificity (Table 1). Slides with primary antibody staining were kept overnight at RT and then washed 3x of 10 min 1x PBS. Secondary antibodies were added to the dilution recommended by the manufacturer’s instructions: goat anti-Ms conjugated to Alexa Fluor 488 (Abcam AB150113, 1:750) and goat anti-Rb conjugated to TRITC (Abcam AB6718, 1:750). Slides with secondary antibody staining were incubated overnight at 4°C and then rinsed 2x of 5 min1x PBS and 1x of 5min 1x PBS with DAPI (1:50,000 dilution). Coverslips were mounted onto slides using cytoseal^TM^ 60.

**Table 1.**
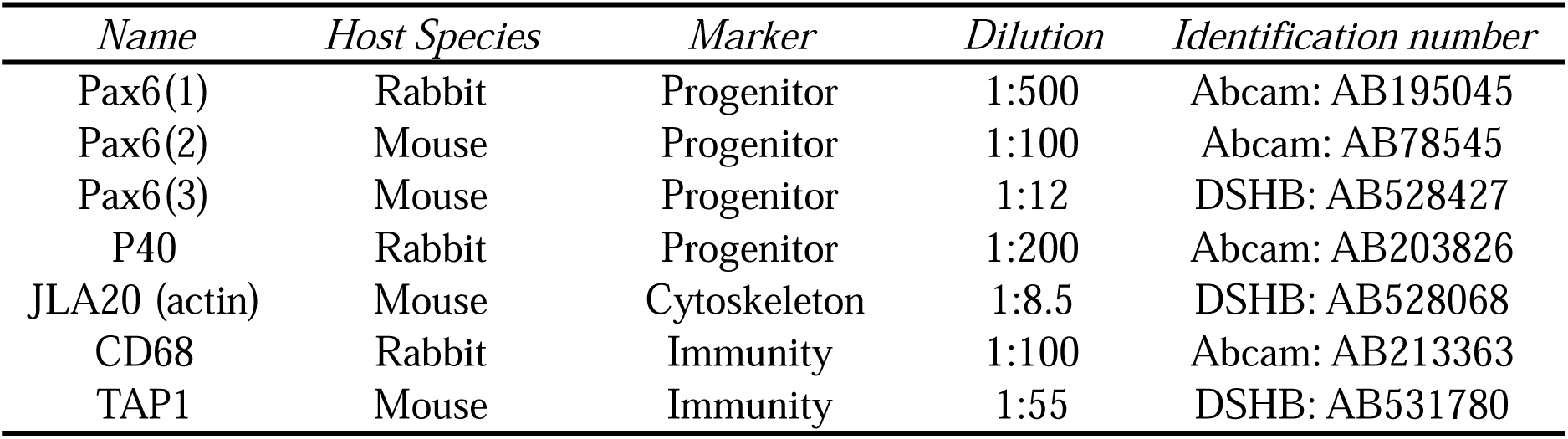
Overview of primary antibodies, their source, and their dilutions used in this study.

### 3.3 Microscopic Imaging and Analysis

Images were taken on a Keyence Fluorescence BZ-X800 microscope. The editing and analysis of images was done in Keyence analyzer software. No primary negative controls for each antibody were used to determine background noise and then those settings were used to obtain final images for each primary antibody staining. Each figure image represents an image that shows the best visualization of that technique. All central cornea and limbal images per technique and antibody came from the same set of tissues. Sample numbers can be found in the figure captions. Figs. 1A,B were made in the tiling and stitching program included in the Keyence analyzer software with a sample number of one each. FIJI image J and Microsoft PowerPoint were used for final image preparation and figure production though no significant changes were made to each image that would change their primary integrity.

**Fig. 1.**
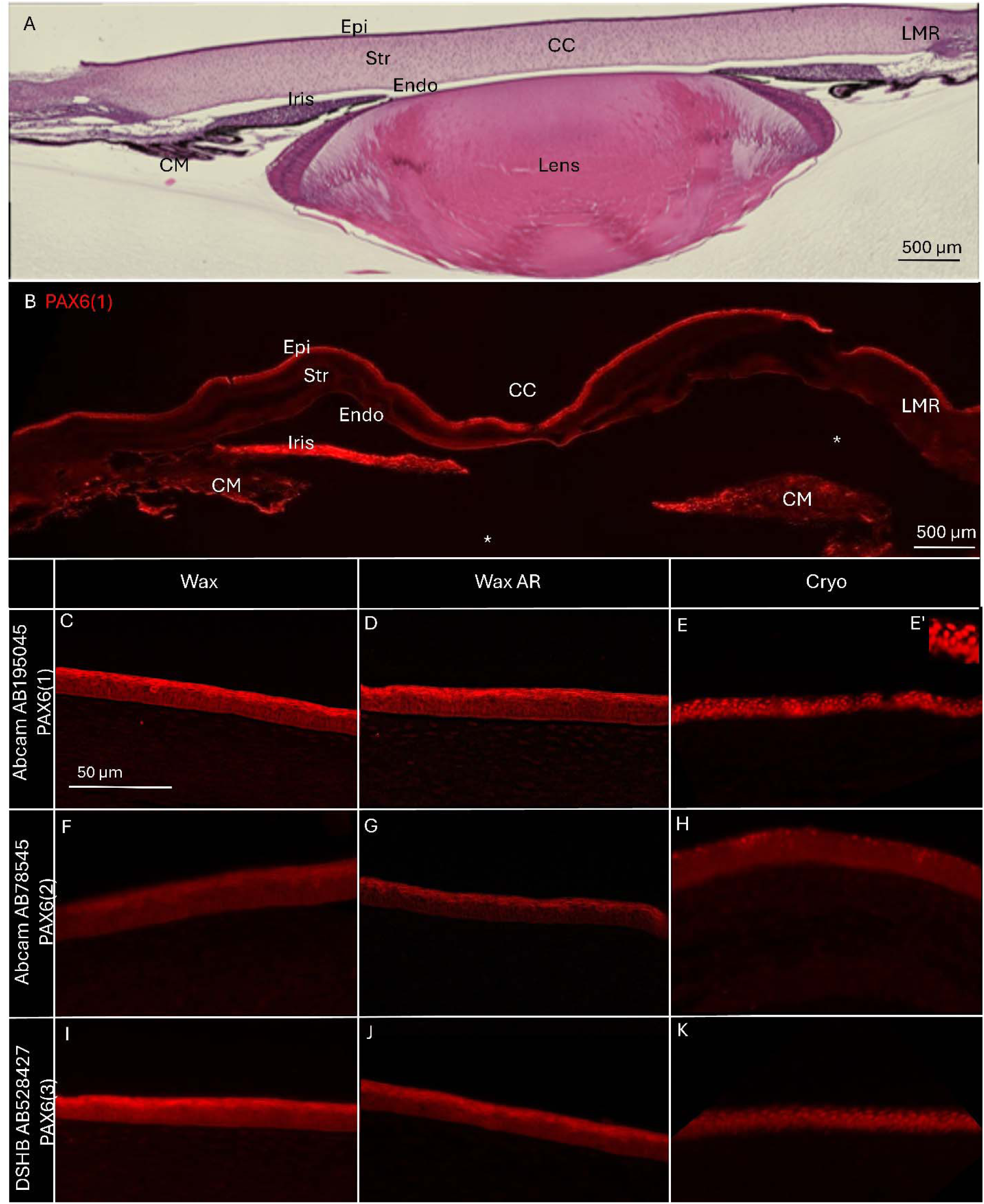
Overview of morphological features by histology and immunohistochemistry of PAX6 of the E18 chicken cornea. (A) Representative H&E-stained stitched section of an E18 chicken eye anterior segment, including the corneal epithelium (Epi), stroma (Str), endothelium (Endo), central cornea (CC), limbal region (LMR), ciliary muscle (CM), iris, and lens. (B) IF stitched section showing PAX6 localization within the E18 chicken eye anterior segment. * represents a missing iris and lens from the section. High-magnification IF images of central corneal sections stained with (C–E, n=10, 8, 12 respectively) anti-PAX6 antibody Abcam AB195045, (F-H, n=8, 9, 13 respectively) Abcam AB78545, and (I-K, n=11, 8, 10 respectively) DSHB AB528427 across different tissue processing methods. (E’) High resolution of the nuclear expression of PAX6 in the epithelium.

Image analysis was performed using the following criteria and results are shown in Table 2. Specificity and signal strength were analyzed by two people for each sample for each antibody and numbers were averaged. Since we were analyzing staining quality across multiple cell types, tissues, and regions we could not use a standard mean fluorescent intensity protocol and instead used a more holistic approach as described. A 0-3 ranking was done for signal strength with the following criteria: 0 (Negative): no staining is observed, or it looks identical to the background; 1 (Weak): barely perceptible or light staining (sometimes described as faint); 2 (Moderate): intermediate staining, clearly visible but not exceptionally bright; 3 (Strong): intense, brightly colored staining. A 0-3 ranking was done for specificity with the following criteria: 0 (Negative): no staining is observed, or it looks identical to the background; 1 (Weak): mostly misexpressed; 2 (Moderate): shows some misexpression and some correct labeling; 3 (Strong): correctly labels cells. If samples included central and limbal regions then both were taken into consideration for an overall number per sample. Overall quality was determined as poor, moderate, or excellent based on the combined analysis of the specificity and signal strength categories (Table 3). Two strong marks resulted in excellent overall quality performance, two moderate marks resulted in moderate overall quality performance, and two weak marks resulted in poor overall quality performance. The CD68 antibody had misexpression in the corneas for all processing techniques and therefore resulted in a poor overall quality ranking even though it had moderate signal strength since marking the correct cells is the most important criteria.

**Table 2.** Resulting marks of averaged antibody specificity and signal strength assessment in corneas.

| *Specificity* |  |  |  | *Signal Strength* |  |  |
| --- | --- | --- | --- | --- | --- | --- |
| *Antibody* | *Wax* | *Wax AR* | *Cryo* | *Wax* | *Wax AR* | *Cryo* |
| Pax6(1) | Weak | Weak | Strong | Strong | Strong | Strong |
| Pax6(2) | Weak | Weak | Moderate | Moderate | Moderate | Moderate |
| Pax6(3) | Weak | Weak | Strong | Strong | Strong | Strong |
| P40 | Negative | Negative | Strong | Negative | Negative | Strong |
| JLA20 (actin) | Moderate | Strong | Moderate | Moderate | Strong | Strong |
| TAP1 | Weak | Weak | Strong | Weak | Moderate | Strong |
| CD68 | Weak | Weak | Weak | Moderate | Moderate | Moderate |

**Table 3.** Overview of antibody quality assessment in corneas based on performance in experimental applications.

| *Antibody* | *Best Processing Technique in Cornea* | *Overall quality* |
| --- | --- | --- |
| Pax6(1) | Cryo | Excellent |
| Pax6(2) | Cryo | Moderate |
| Pax6(3) | Cryo | Excellent |
| P40 | Cryo | Excellent |
| JLA20 (actin) | Wax AR | Excellent |
| CD68 | none | Poor |
| TAP1 | Cryo | Excellent |

### 3.4 Results and Discussion

An overview of the results of different IHC processing methods in embryonic chicken corneal tissue can be found in Table 3. Summarily, all the PAX6 antibodies, the P40 antibody, and the TAP1 antibody worked best with the cryosectioning technique. Actin gave the best results with wax AR and CD68 showed limited or no corneal expression across all techniques while showing positive expression in the kidney across all techniques. Fig. 1A shows the expected normal histology of a 30 image stitched H&E stain of the anterior region of the E18 chicken eye for comparative purposes. Fig. 1B shows a stitched picture of 30 images of a cryosectioned anterior region of an E18 chicken eye, which includes PAX6(1) staining of the epithelium of the central cornea, limbal region, iris, and ciliary muscle.

#### 3.4.1 Progenitor activity

PAX6 is an important transcription factor involved in eye development, stem cell regulation, and corneal cellular maintenance. We tested three PAX6 antibodies from two companies, Abcam and DSHB, for immunofluorescence (IF) on the E18 chicken central corneal epithelium using three different processing techniques: wax-embedding, wax AR, and cryosectioning (Fig. 1). Our results show that only the cryosectioning technique gave nuclear-specific epithelial-only staining, and the wax-embedded and wax AR processing techniques gave non-specific (non-nuclear) staining, showing higher signal at the epithelial superficial layer, which is not expected (Fig. 1C,D,F,G,I,J). Furthermore, none of the antibodies showed expression in the central cornea stromal layer as expected.

The two Abcam antibodies are considered suitable for human, mouse, and rat for applications such as Western blot (WB), IHC, and IF by the company’s datasheet. However, they have never been tested on chicken or corneas before. Our data show that both the Abcam antibodies did work with cryosectioning though with slightly different results. PAX6(1) showed strong specificity and strong signal with expression found in the nucleus of all cells in the epithelium and no expression in the stroma, as expected (Fig. 1E,E”). Fig. 1H shows PAX6(2) also worked only in the epithelium and not the stroma but with moderate signal strength and specificity. Some of the more superficial cells had more specific nuclear expression, but the signal was not consistent throughout all the layers of the epithelium or across the whole central region (Fig. 1H). The exact immunogen information is considered propriety information, so it is unknown why these two antibodies would give different results for the cryosectioning technique.

The PAX6(3) antibody purchased from DSHB is stated by the company to be used for WB, IHC, IF, FACS, and ChIP applications. DSHB also states that the antigen species is chicken and the antibody is from the N-terminal, aa 1-223. It has been published with successful IF staining in chicken brain and retina, and mouse iris, as well as a myriad of other species and organs (Peretz et al, 2016; Zhang et al., 2016; Hagglund et al., 2017). Furthermore, the referenced experiments were done with cryosectioning or by using a whole-mount with no sectioning necessary. Therefore, our results complement these previous experiments, showing that cryosectioning is best for this antibody across organs and species and is successful for chicken corneas (Fig. 1K).

The monoclonal P40 – DeltaNp63 antibody from Abcam binds to the P40 protein that acts as a transcription factor in regulating epithelial stem cell maintenance and differentiation, and its nuclear expression is usually found in the basal layer of the skin and select organs, including the cornea. Its recommended applications by Abcam are IF, IHC, immunoprecipitation (IP), WB, and flow cytometry. We tested this antibody for IF on the E18 chicken cornea in the epithelium, stroma, and endothelium of the central and limbal regions using three different processing techniques: wax-embedding, wax AR, and cryosectioning (Fig. 2).

**Fig. 2.**
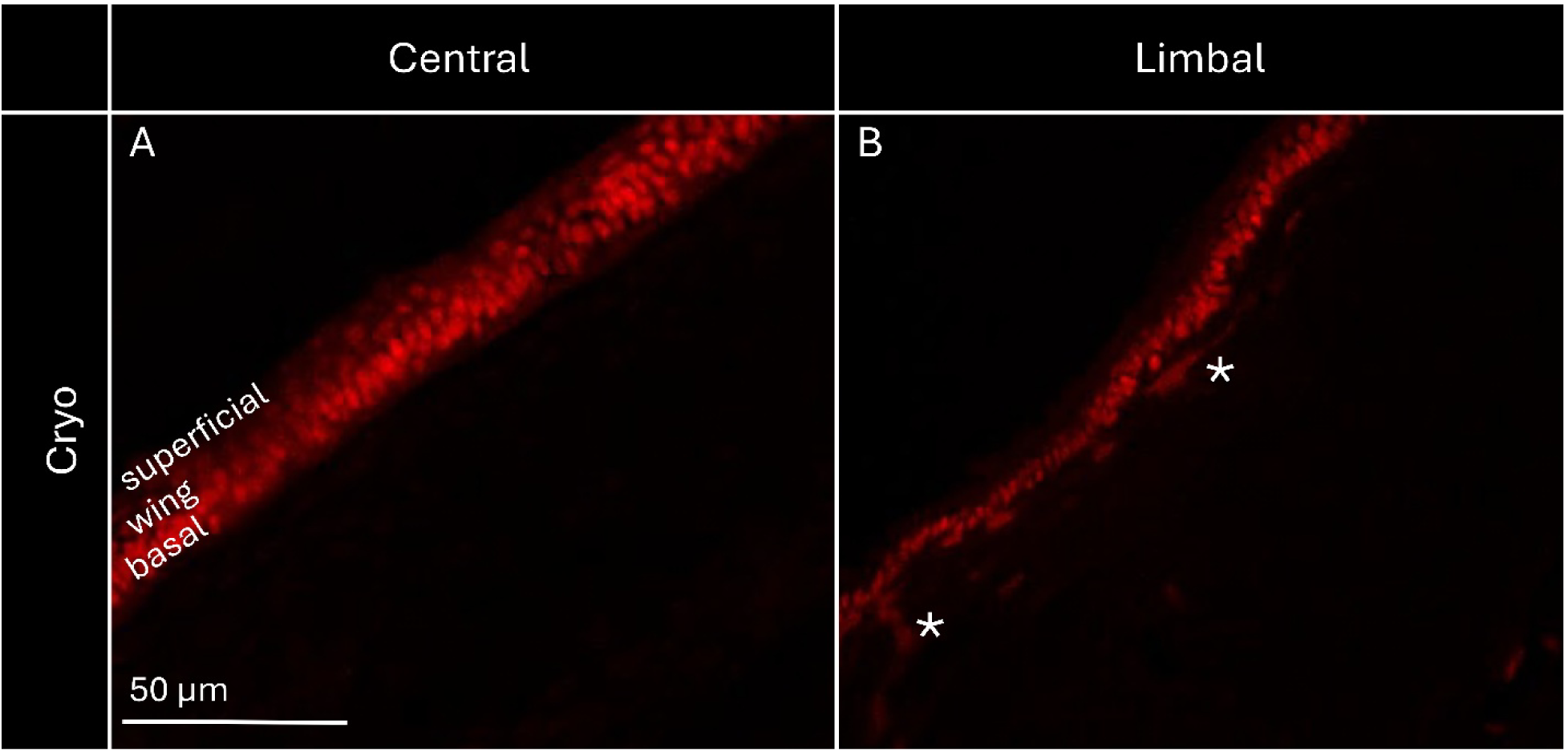
P40 IF staining in E18 chicken cornea highlighting epithelial progenitor populations. Representative images show the central cornea (A, n=9) and limbal region (B, n=9). Regional distribution of cells in the epithelium is highlighted by the superficial, wing, and basal markings. * represents P40 expression seen in the stroma.

Our results show the gradient of P40 expression across epithelial layers as expected with strong nuclear expression in the basal cell layer, moderate expression in the suprabasal (wing) cells, and weak expression in the superficial cells of the corneal epithelium throughout the central and limbal regions (Fig. 2). Some scattered P40 expression can be seen in the stroma just below the epithelium for the limbal region (Fig. 2B), which was unexpected. The corneal stromal stem cells are mesenchymal-like stem cells that have the potential to differentiate into keratocytes, support the limbal epithelial stem cells, and aid in corneal stroma regeneration. Though our data shows these cells are P40 positive, more work needs to be done to see if these cells coexpress with other corneal stromal stem cell markers. Additionally, our experiments revealed that the antibody only worked using cryosectioning with negative results when processed with wax (n=4) and wax AR (n=5) techniques (data not shown). No expression was seen in the endothelial layer.

#### 3.4.2 Tissue Architecture

The monoclonal antibody JLA20 from DSHB binds the chicken antigen actin, a cytoskeletal protein. This antibody recognizes all isoforms of actin over a broad range of species, and its recommended application by DSHB is for IHC, IP, and WB. Actin is a highly prevalent protein found in epithelial cells, keratinocytes of the stroma, and endothelial cells, which make up the most internal layer of the cornea and the blood vessels found in the limbal regions, and therefore, it should show expression in all layers of the cornea. However, the mature collagen fibrils of the stroma that help provide the main optical function of the cornea do not contain actin and should not show expression of JLA20. Actin is upregulated in wound healing for the stroma (Ritchey et al., 2011), and having a dependable chicken-specific actin antibody for corneal wound healing experiments is important for the biomedical field. We tested this antibody for IF on the E18 chicken cornea in the epithelium, stroma, and endothelium of the central and limbal regions using three different processing techniques: wax-embedding, wax AR, and cryosectioning (Fig. 3).

**Fig. 3.**
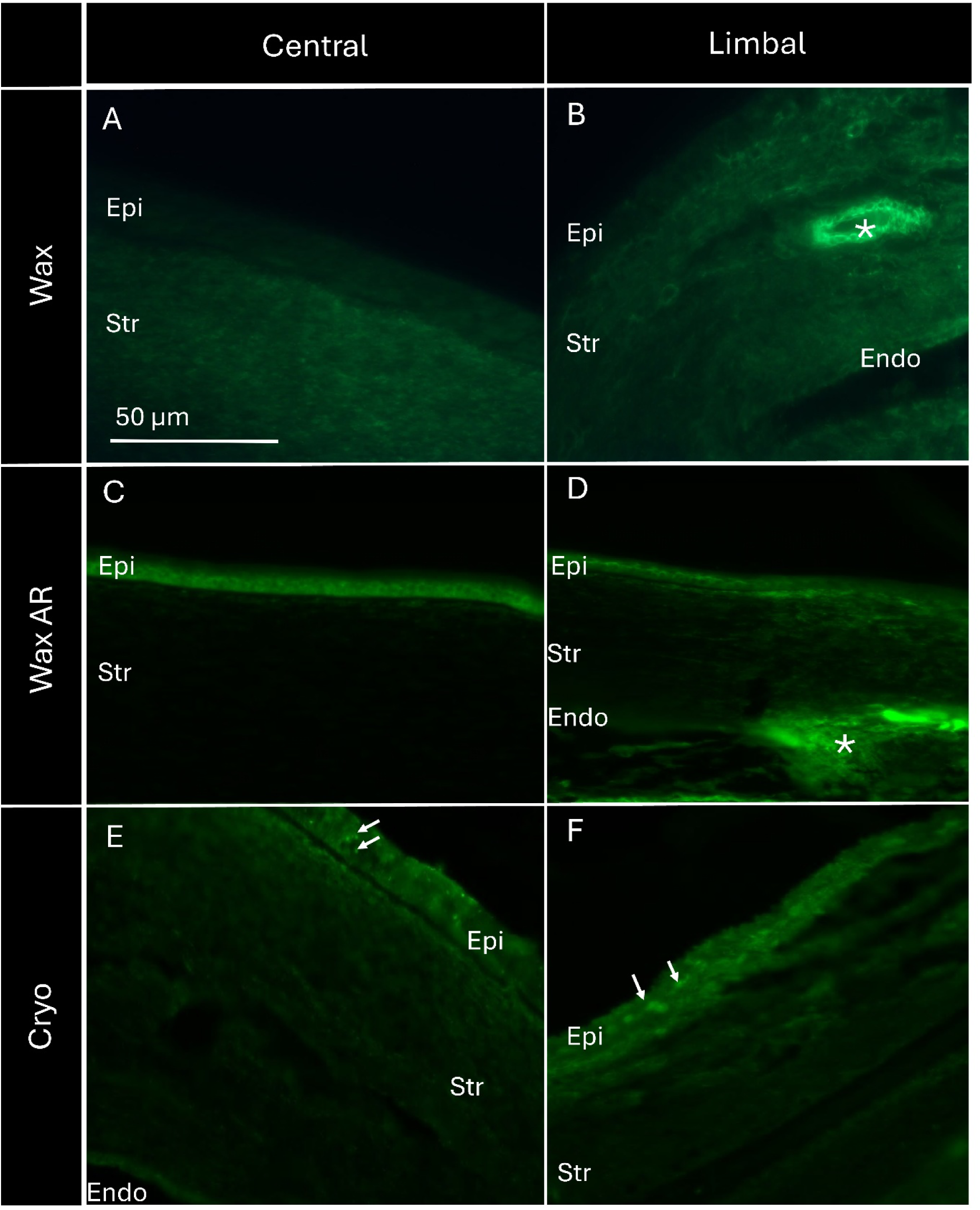
IF staining for JLA20 (actin) in E18 chicken corneal sections, comparing central and limbal regions across different embedding methods. Sections were processed using wax embedding (A–B, n=7), wax AR (C–D, n=9), or cryosectioning (E–F, n=8). Corneal epithelium (Epi), stroma (Str), endothelium (Endo), * marks blood vessels, and arrows indicate punctated staining.

The staining revealed the antibody worked across tissues and regions of the cornea but with differences in signal strength and specificity depending on the processing method. Our results show better overall quality in wax AR and cryosectioning processing methods compared to standard wax embedding (Fig. 3). However, the cryosectioning process also resulted in brightly punctated spots in the epithelium that should not be present (Fig. 3E,F). The wax-embedded stains showed weaker expression in the epithelium and endothelium compared to the stroma, while wax AR and cryosectioning showed stronger expression in the epithelium and endothelium compared to the stroma in corneal regions, which is the expected result. Actin is also highly expressed in blood vessels (Fig. 3B,D) found in the limbal region and shows strong signal staining for the wax and wax AR methods. Together, this information results in wax AR as the best processing method for the cornea to show actin in the central cornea and limbal regions.

#### 3.4.3 Immune surveillance

The monoclonal antibody TAP1 from DSHB binds chicken class I MHC antigens primarily found on B cells, macrophages, and dendritic cells. DSHB recommends its application for IF, IHC, and IP. The monoclonal antibody CD68 from Abcam binds human CD68 antigens, a transmembrane glycoprotein primarily used as a pan-macrophage marker for identifying monocytes, macrophages, and dendritic cells. Abcam recommends its application for IF, IHC, and WB. Though both antibodies bind immune cells, such as macrophages and dendritic cells, they bind different antigens. We tested these antibodies for IF on the E18 chicken cornea and the E18 chicken kidney as an experimental control using three different processing techniques: wax-embedding, wax AR, and cryosectioning (Figs. 4,5).

**Fig. 4.**
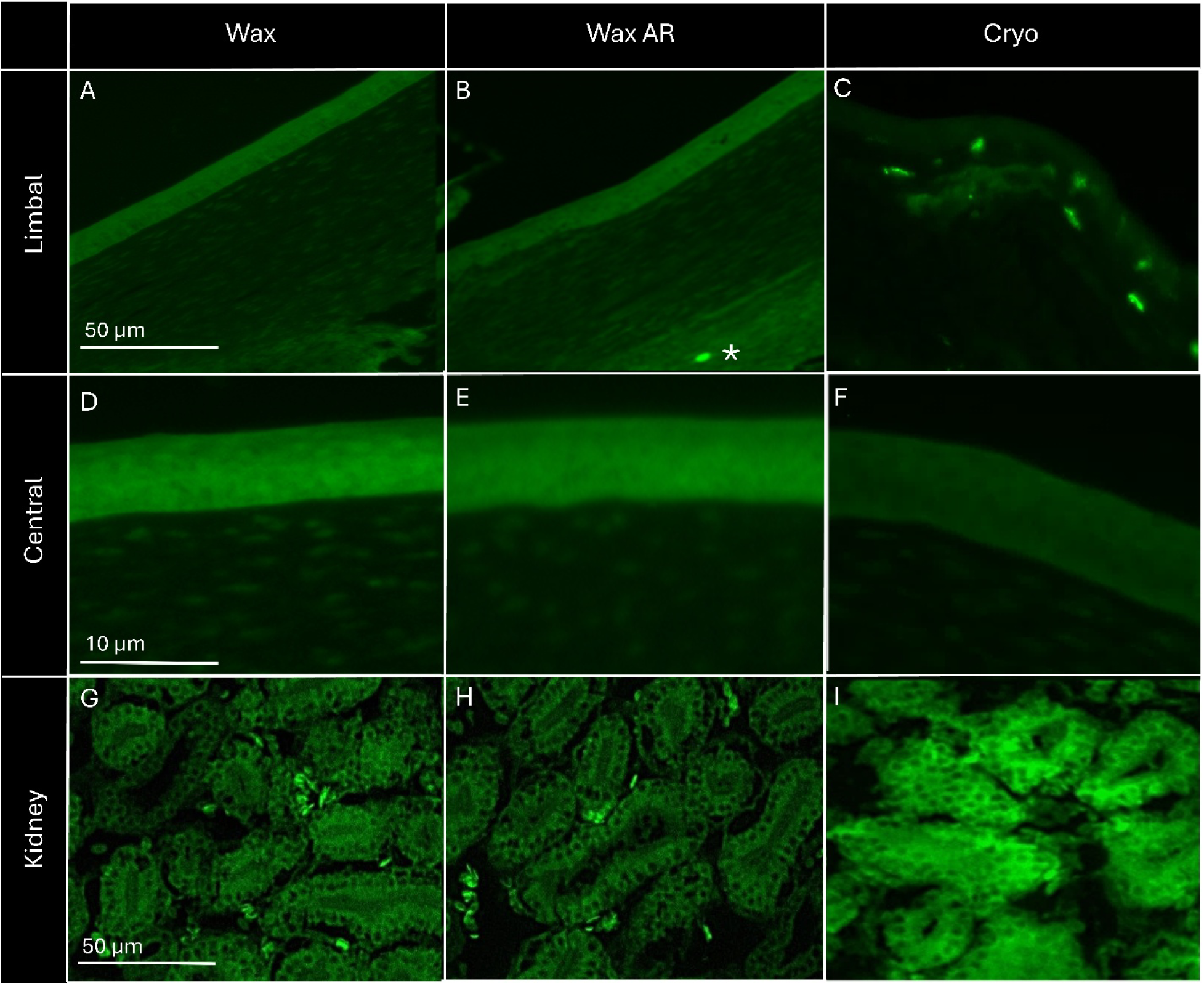
E18 chicken corneas and kidneys comparing TAP1 IF across wax-embedded, wax AR, and cryosectioned tissues. (A–C) Limbal region, showing TAP1 signal primarily associated with the stromal compartment, with marked differences in signal intensity and continuity across processing methods. (D–F) Central cornea, showing background or negative staining across all processing methods. (G–I) Kidney tissue, serving as a TAP1-positive control, showing robust TAP1 expression with clear cellular and subcellular definition in wax and wax AR sections and reduced structural resolution in cryosectioned tissue. Sample numbers for eye tissues used in the techniques of wax, wax AR, and cryo were respectively 8, 10, and 14 and 2 kidneys were used per technique. * marks one positive cell in stroma

**Fig. 5.**
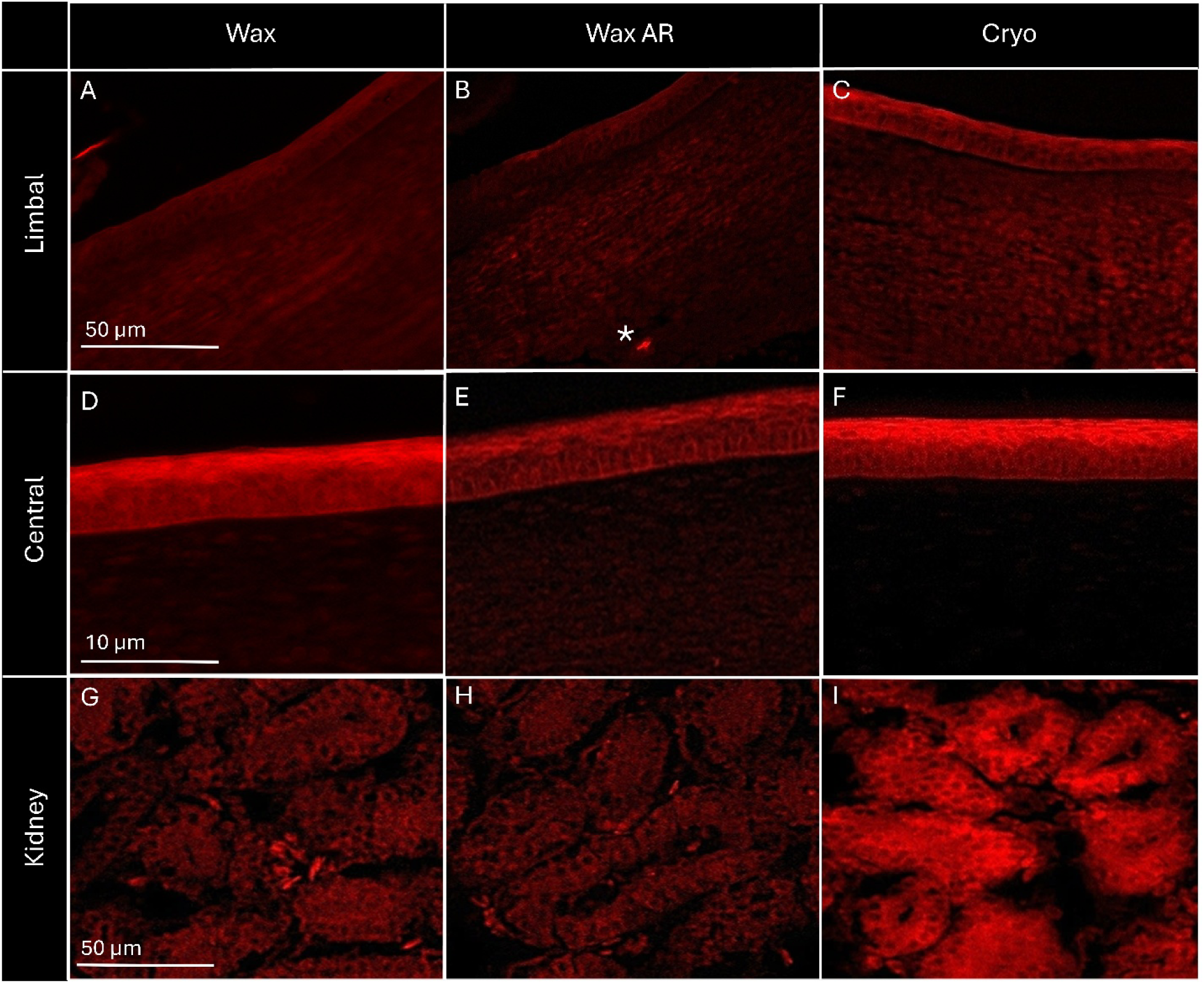
E18 chicken corneas and kidneys comparing CD68 IF across wax-embedded, wax AR, and cryosectioned tissues. (A–C) Limbal region, showing mostly negative and background CD68 signal across processing methods. (D–F) Central cornea, showing background or negative staining across all processing methods. (G–I) Kidney tissue, serving as a CD68-positive control, showing robust CD68 expression with clear cellular and subcellular definition in wax and wax AR sections and reduced structural resolution in cryosectioned tissue. Sample numbers for eye tissues used in the techniques of wax, wax AR, and cryo were respectively 6, 5, and 8 and 2 kidneys were used per technique. * marks one positive cell in stroma

The staining showed TAP1 and CD68 worked in the chicken kidney for all three processing techniques (Fig. 4,5G-I) as expected when compared to the Edgtton et al. (2008) paper that examined the phenotype of dendritic cells in the mouse renal interstitium after intrarenal ovalbumin injection. Their research used CD11c as a marker for dendritic cells; however, the expression pattern should be similar for the chicken kidney with TAP1 and CD68, since all three proteins are markers for dendritic cells. The kidney was used as a positive control since resident immune cell numbers are low in the central and limbal corneal regions, making false negatives a potential result (Hamrah et al., 2003). Since the two antibodies have different host species we were able to use the same tissue for both antibodies.

Macrophages and dendritic cells are known to be present in the stroma of the limbal region and present in much lower numbers in the central cornea and the epithelium of the limbal region (Niederkorn, 2002; Hamrah et al., 2003; Prieto-Nevarez et al., 2026). Our results showed that cryo-preserved tissue had TAP1-positive cells in the stroma of the limbal region and no positive cells in the limbal epithelium or the central cornea (Fig. 4C,F). The wax processing techniques showed mostly negative results for all corneal tissue and high background staining for the central cornea (Fig. 4A,B,D,E). Figure 4B shows one positive cell in the limbal stroma, but one positive cell is not the expectation for that region as multiple cells are expected as seen in the cryo-preserved tissue (Fig. 4C). CD68 showed mostly negative results with high background for all the techniques and tissues, and potential misexpression at the corneal edge in the wax method (Fig. 5). Figure 5B does show the same one positive cell in the wax AR technique that showed positive for TAP1 in Figure 4B. The same tissue was used with double staining, TAP1 and CD68.

## 4. Potential Pitfalls and Troubleshooting

With any IHC the most common pitfalls are improper tissue fixation, high background noise, and potential incorrect antibody staining due to incorrect concentrations. Considering this is a research paper about diverse types of tissue fixation across seven different antibodies tested in a heterogenous new tissue, the chicken cornea, we tried to compensate for these common IHC issues as much as possible. We controlled experimental components like fixation time (24hrs), fixation solution (4% PFA), buffers, and dehydration techniques so that the focus was on the differing fixation processing methods (wax, wax AR, or cryo). Therefore, this experimental approach was specifically set to explore the different tissue fixation techniques while keeping all other experimental components the same for each of the antibodies. Due to the nature of the ocular environment and the presence of various proteins, there is a significant risk of non-specific immunofluorescent staining. Using positive and negative controls for each tissue and staining helped to recognize and eliminate background noise and autofluorescence prevalent in IHC staining. All experimental images were taken at the settings for the negative control. To compensate for too much or too little primary or secondary antibody we used the manufacturer’s concentration suggestion with the positive control tissue first and then optimized for best results. All experimental procedures were completed with the optimized concentrations found in Table 1. Since CD68 antibody did not work in the cornea with any of the fixation techniques but did in the positive control kidney tissue that was processed at the same time more experiments would need to be done with the other procedural components however this was beyond the scope of this work.

Due to the heterogeneity of tissue structure and markers we were unable to use a standard mean fluorescent intensity protocol but instead used a more global approach to antibody overall quality, including specificity and signal strength. This qualitative approach was successful for this study, however using a more quantitative analysis would be useful for a smaller, more specific study for each antibody.

It is common for cryopreserved tissue to show blurrier or lower-resolution morphology when compared to wax-embedded techniques, and this phenomenon explains the microscopic resolution difference between the wax- and cryo-embedded techniques. Wax-embedded tissue usually is the superior method for fine cellular architecture preservation, something that is important when looking at small populations of cells within complex tissues. To get the best resolution of the cells we used the haze reduction tool on the Keyence BZ-X800 microscope, but a confocal microscope would provide higher clarity.

We picked this set of markers partly due to the convenience of access but also because they were within the range of attributes used to research corneal wound healing, a necessary field in perpetuating corneal health. We decided to focus on PAX6 due to its significance in corneal health such as its involvement in Aniridia and corneal regeneration. There are other markers not used in this study that can differentiate tissue regions or composition like cytokeratin for central-specific epithelial cells or ABCG2 as a surface marker for stem cells, but we limited our selection due to availability of time, money, and interest level. We believe the current selection encompasses a good range of markers for corneal structure and the basics of wound healing.

## 5. Conclusion

In this methodological research, we compared three IHC processing methods with seven antibodies of five different proteins in embryonic chicken corneal tissue. We assessed the overall imaging quality and cellular structural integrity of IF staining with the three processing methods, concluding an overall best processing technique for each antibody across the different tissues within the embryonic chicken cornea (Table 3). Overall, the three PAX6 antibodies, the P40 antibody, and the TAP1 antibody had the best quality assessment with cryosectioning; actin was preserved best with wax AR; and CD68 did not work in the cornea but did in the kidney positive control.

In conclusion, we recommend using cryosectioning to best preserve nuclear antigens like PAX6, P40, and TAP1 in the epithelium and stroma of the embryonic chicken cornea. However, actin, an example of tissue architecture, is best preserved with wax AR, though can be successful with the other processing techniques. Ultimately, this study provides a practical reference for future studies that use immunostaining of progenitor activity, tissue architecture, and immune surveillance in experiments for chicken eyes.

## Author statement

All authors have seen and approved the final version of the manuscript being submitted and warrant that the article represents the authors’ original work, has not received prior publication and is not under consideration for publication elsewhere.

## Acknowledgements

We would like to thank Tyson Farms for the fertile chicken eggs.

## Declaration of generative AI and AI-assisted technologies in the manuscript preparation process

During the preparation of this work the author(s) used Grammarly in order to provide grammar and edit suggestions and ChatGPT for reference styling. After using the service, the authors reviewed and edited the content as needed and take full responsibility for the content of the published article.

